# *APOE* deficiency impacts neural differentiation and cholesterol biosynthesis in human iPSC-derived cerebral organoids

**DOI:** 10.1101/2022.06.30.498241

**Authors:** Jing Zhao, Tadafumi C. Ikezu, Wenyan Lu, Jesse R. Macyczko, Yonghe Li, Laura J. Lewis-Tuffin, Yuka A. Martens, Yingxue Ren, Yiyang Zhu, Yan W. Asmann, Nilüfer Ertekin-Taner, Takahisa Kanekiyo, Guojun Bu

## Abstract

The apolipoprotein E (*APOE*) gene is the strongest genetic risk factor for Alzheimer’s disease (AD); however, how it modulates brain homeostasis is not clear. The apoE protein is a major lipid carrier in the brain transporting lipids such as cholesterol among different brain cell types. Here, we show that *APOE* deficiency in human iPSC-derived cerebral organoids impacts brain lipid homeostasis by modulating multiple cellular and molecular pathways. Molecular profiling through single cell RNA-sequencing revealed that *APOE* deficiency leads to changes in cellular composition of isogenic cerebral organoids likely by modulating the EIF2 signaling pathway as these events were alleviated by the treatment of a pathway inhibitor ISRIB. *APOE* deletion also leads to activation of the Wnt/β-catenin signaling pathway with concomitant decrease of *SFRP1* expression in glia cells. Importantly, the critical role of apoE in cell type-specific lipid homeostasis was observed upon *APOE* deletion in cerebral organoids with a specific upregulation of cholesterol biosynthesis in excitatory neurons and excessive lipid accumulation in astrocytes. Relevant to human AD, *APOE4* cerebral organoids show altered neurogenesis and cholesterol metabolism compared to those with *APOE3*. Our work demonstrates critical roles of apoE in brain homeostasis and offers critical insights into the *APOE4*-related pathogenic mechanisms.

## Introduction

The dysregulation of lipids has emerged as a key feature of several age-related neurodegenerative diseases (1–4). Apolipoprotein E (apoE) is a major component of brain-derived lipoproteins with HDL-like properties. ApoE is produced primarily by astrocytes in the brain and facilitates the transfer of cholesterol and phospholipids between cells in the brain (5–7). Astrocytes metabolically interact with neighboring neurons by providing cholesterol, phospholipids, hydrophobic vitamins, and antioxidants (8–10). Indeed, astrocytic apoE has been associated with multiple neuronal functions, including axon guidance, survival, amyloid-β (Aβ) metabolism, neurogenesis, and synaptic plasticity (5, 6, 11). Since apoE also plays a critical role in lipid efflux from brain cells (12), apoE is predicted to mediate diverse functions in both cell-autonomous and non-cell-autonomous manners (5, 9, 13–15). In humans, the *APOE* gene exists as three polymorphic alleles (*APOE2*, *APOE3*, and *APOE4*), where *APOE4* is the strongest genetic risk factor for Alzheimer’s disease (AD) (16–18). *APOE4* has been shown to contribute to AD pathogenesis through multiple Aβ-dependent and independent pathways including lipid transport and metabolism (6, 19–21). While *APOE4* increases tau phosphorylation and Aβ production in human induced pluripotent stem cell (iPSC)-derived neurons (14, 15), iPSC-derived astrocytes carrying *APOE4* show the exacerbated cholesterol/lipid droplet accumulation, diminished cholesterol secretion, and impaired neurotrophic functions compared to those with *APOE3* (22–24). Thus, better understanding of how apoE regulates the crosstalk between different brain cell types is critical to understanding how it modulates brain homeostasis and AD risk and facilitate rational design of apoE-targeted treatment strategies against AD (25, 26).

With the development of iPSC technologies, the emergence of three-dimensional cerebral organoid model system with distinct cell diversity provides an optimal tool to define the cell-type-specific effects of disease associated genes in their native environment (27–29). Thus, to dissect the roles of apoE in different brain cell types, we conducted single-cell RNA sequencing (scRNA-seq) analysis and pathway validation in the cerebral organoids from *APOE* deficient (*APOE*^-/-^) and isogenic parental iPSC lines. We found that *APOE* deficiency impacts cellular composition of the iPSC-derived cerebral organoids by modulating the EIF2 pathway and cholesterol biosynthesis pathway. Some of our observations were further extended to *APOE4* cerebral organoids. Our findings provide new insights into the cellular and molecular mechanisms through which apoE modulates brain homeostasis in a cell-type-specific manner.

## Results

### Cellular diversity of human iPSC-derived cerebral organoids revealed by scRNA-seq

We generated three-dimensional (3-D) cerebral organoids from a human parental iPSC line (Xcell Science) and its isogenic *APOE* deficient (*APOE^-/-^*) iPSC line (Xcell Science) following our previously reported protocol (30, 31). To elucidate the cell type-specific effects of *APOE* deficiency in the cerebral organoids, we performed Gel Bead-In-EMulsion (GEM)-based scRNA-seq in the parental and *APOE^-/-^* cerebral organoids at Day 90 (**Supplementary Figure 1a**). The cerebral organoids were enzymatically dissociated and analyzed by scRNA-seq (14920 cells from control cerebral organoids and 12279 cells from *APOE^-/-^* cerebral organoids) after quality control filtering.

All scRNA-seq datasets were aggregated and analyzed following the standard Seurat package procedures (v.4), and cells were clustered based on their expression of variable genes. The analysis by t-distributed stochastic neighbor embedding (t-SNE) revealed 19 transcriptionally distinct clusters (**Figure 1a**), where clusters were classified to 5 major cell types (**Figure 1a, b, Supplementary Figure 1b-i**) according to the expression of known cell type-specific markers (29, 32–34); excitatory neurons (ExN, cluster 0, 1, 2, 3, 5); radial glia (RG, cluster 4, 7, 9, 11, 17); astrocytes (Astro, cluster 10, 12, 13); intermediate progenitor cells (IPC, cluster 6) and inhibitory neurons (InN, cluster 8, 14, 15). Two clusters were defined as undecided clusters (UD, cluster 16, 18) due to the lack of distinct marker expression. Notably, within the excitatory neuron and astrocyte population, the expression of specific marker genes in different clusters showed a pattern of neuronal layer differentiation and astrocyte maturation (**Figure 1b**). Together, these results illustrate the cell type diversity within the iPSC-derived cerebral organoids.

**Figure 1.**
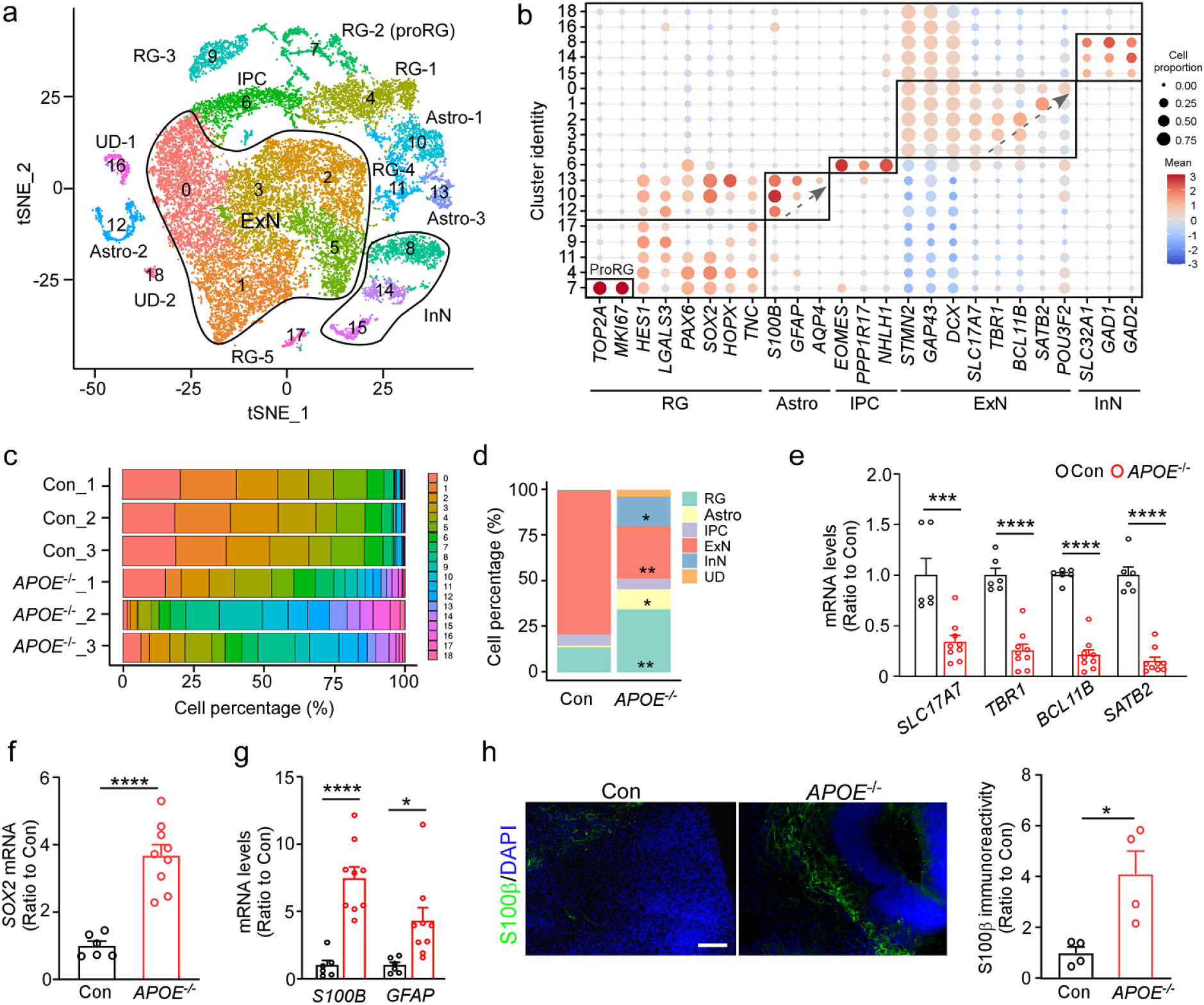
*APOE* deficiency leads to cellular composition changes in iPSC-derived cerebral organoids. **a-d** Parental control and isogenic *APOE^-/-^* iPSCs were differentiated into cerebral organoids and dissociated into single cells at Day 90 for scRNA-seq analysis. Three cerebral organoids were pooled and analyzed as one sample (n=3 samples/genotype). (a) t-SNE plot and cluster identification of scRNA-seq data. Cell clusters were defined based on the expression of cluster marker genes and known marker genes. ExN, excitatory neuron; RG, radial glia; InN, inhibitory neuron; IPC, intermediate progenitor cell; Astro, astrocyte; UD, undecided. (b) Dot plot of canonical genes to classify tSNE clusters. Cluster identities are labeled on the left and canonical marker genes are indicated on the bottom. (c) Percentage of cell clusters in each cerebral organoid sample. (d) Comparison of the percentage of different cell types between control and *APOE^-/-^* cerebral organoids. **e-g** The mRNA levels of cerebral layer markers (e; *SLC17A7*, *TBR1*, *BCL11B* and *SATB2*), neural stem cell marker (f; *SOX2*) and astrocytic markers (g; *S100B* and *GFAP*) were quantified by RT-qPCR. Three cerebral organoids were pooled and analyzed as one sample (n=6-9 samples/genotype). **h** Astrocyte differentiation in cerebral organoids were evaluated by S100β immunostaining (n=4 organoids/genotype). Scale bar: 50 μm. Experiments were repeated in two independently differentiated batches (e-h). All data are expressed as mean ± SEM. Student’s t tests were performed to determine statistical significance. *p<0.05, **p<0.01, *** p<0.001, ****p<0.0001.

In the brain, apoE is manly produced by astrocytes, activated microglia, vascular mural cells and choroid plexus cells, and to a lesser extent by stressed neurons (6). To evaluate the cell type-specific *APOE* expression in the cerebral organoids, *APOE* gene expression levels in different cell clusters were plotted based on the scRNA-seq data (**Supplementary Figure 2**). In the parental control cerebral organoids, *APOE* is predominantly expressed in radial glia and astrocyte populations, with subtle expression in the neuronal population (**Supplementary Figure 2a)**, while apoE immunostaining showed relative diffuse distribution (**Supplementary Figure 2b**). These results indicate that glia cells are the main source of apoE production in the iPSC organoids. *APOE* expression was not detected in any cluster of *APOE^-/-^* cerebral organoids, immunostaining showed background apoE expression in *APOE^-/-^*cerebral organoids.

### Altered cellular composition in *APOE* deficient cerebral organoids

The cerebral organoids differentiated from iPSCs have been found to resemble the developing brain and exhibit different cerebral layers expressing specific markers (35, 36). While investigating the effects of *APOE* deficiency on the cellular composition of cerebral organoids, we found dramatic differences between parental and *APOE^-/-^* cerebral organoids (**Figure 1c**). Specifically, excitatory neuron population was significantly reduced by *APOE* deficiency, whereas radial glia, astrocyte and inhibitory neuron populations were increased (**Figure 1d**). To validate the cellular composition changes in the *APOE^-/-^* cerebral organoids, the expression levels of cell type-specific markers were evaluated via RT-qPCR and immunostaining. Consistent with the scRNA-seq data, the mRNA levels of excitatory neuronal marker (*SLC17A7*), deep layer neuron markers (*TBR1* and *BCL11B/CTIP2*) and upper layer neuron marker (*SATB2*) decreased in the *APOE^-/-^* cerebral organoids (**Figure 1e**), indicating that *APOE* deficiency inhibits neuronal layer formation in cerebral organoids. On the other hand, the marker for ventricular zone (*SOX2*) and astrocytic markers *S100B* and *GFAP* increased significantly (**Figure f-g, Supplementary Figure 3a**). Immunostaining also showed a significant increase in the ratio of S100β-positive astrocytes in the *APOE^-/-^* cerebral organoids (**Figure 1f**). Pseudotime trajectory analysis also revealed distinct differentiation trajectories between parental and *APOE^-/-^* cerebral organoids (**Supplementary Figure 3b**). Together, these results suggest an important role of apoE in neural fate regulation in the iPSC-derived cerebral organoids.

### Impaired neural differentiation in *APOE* deficient cerebral organoids associated with activated EIF2 signaling pathway

To investigate how *APOE* deficiency affects gene expression and pathways in different cell types within the cerebral organoids, the differentially expressed genes (DEGs) in the main cell clusters were identified and subjected to the pathway analysis (**Figure 2a-h, Supplementary Figure 4**). The two major cell populations, excitatory neuron and radial glia clusters, showed an overwhelming upregulation of cellular stress-related pathways, including “*EIF2 signaling pathway*” and “*mTOR signaling pathway*” (**Figure 2b, f**). Consistently, Western blotting confirmed the significant increase of eukaryotic initiation factor 2α (eIF2α) phosphorylation in *APOE^-/-^* cerebral organoids compared to the controls (**Figure 2i**). To further investigate the contribution of EIF2 signaling pathway to the phenotypic changes, *APOE^-/-^* cerebral organoids were treated with the integrated stress response inhibitor (ISRIB) which blocks the phosphorylation of eIF2α in the integrated stress response (37) from Day 60 for 30 days. While Western blotting showed a significant reduction of eIF2α phosphorylation by ISRIB in the cerebral organoids (**Figure 2j**), the ISRIB administration restored the altered mRNA levels of neuronal markers (*DCX*, *SLC17A7* and *TBR1*) and astrocytic markers (*S100B* and *GFAP*) in *APOE^-/-^* cerebral organoids (**Figure 2k**). Immunostaining confirmed that ISRIB administration increased CTIP2-positive neurons in the cerebral organoids (**Figure 2l**). Together, these results indicate that *APOE* deficiency leads to altered neuronal differentiation in the cerebral organoids by activating EIF2 signaling pathway.

**Figure 2.**
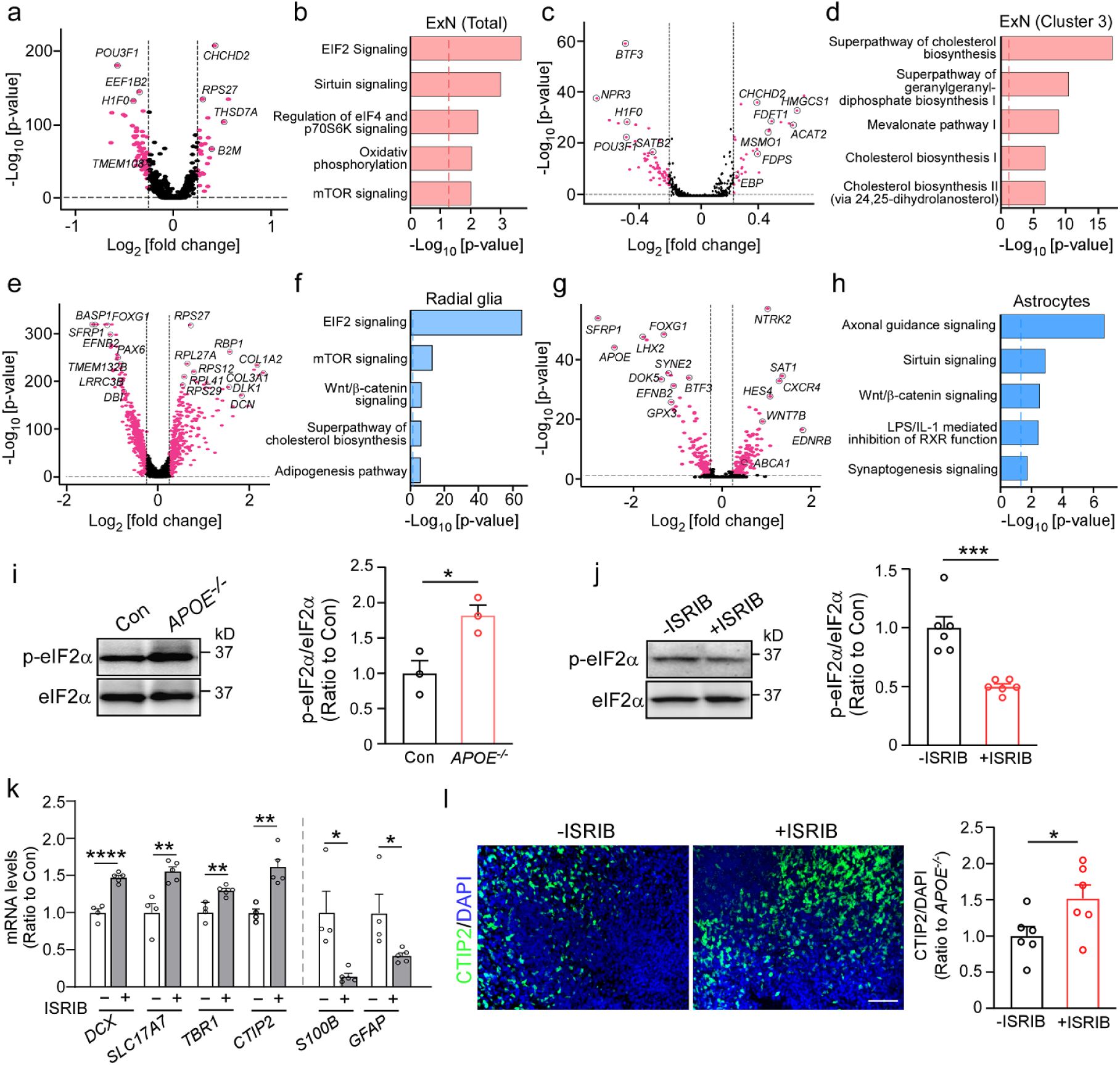
*APOE* deficiency alters neural differentiation in cerebral organoids by activating the EIF2 signaling pathway. **a-h** DEGs in cell clusters were identified through the scRNA-seq and subjected to the Ingenuity pathway analysis. Volcano plot and gene ontology analyses for DEGs (*APOE^-/-^ vs.* Con) in total excitatory neuron clusters (a, b), excitatory neuron cluster 3 (c, d), radial glia clusters (e, f) and astrocyte clusters (g, h) are shown. Genes significant at the P value ≤ 0.05 and fold change ≥ 1.2 are denoted in purple. Selective DEGs are labelled in the volcano plots. **i** Phosphorylation levels of eIF2α in the control and *APOE^-/-^* cerebral organoids at Day 90 were quantified by Western blotting. Three cerebral organoids were pooled and analyzed as one sample (n=3 samples/genotype). **j** *APOE^-/-^* cerebral organoids at Day 60 were treated with or without ISRIB (100 nM) for 30 days. Phosphorylation level of eIF2a in the cerebral organoids was quantified by Western blotting. Three cerebral organoids were pooled and analyzed as one sample (n=6 samples/genotype). **k** The mRNA levels of cerebral layer markers (*DCX*, *SLC17A7*, *TBR1* and *CTIP2*) and astrocytic markers (*S100B* and *GFAP*) were quantified by RT-qPCR. Three cerebral organoids were pooled and analyzed as one sample. All data are expressed as mean ± SEM (n=4-5 samples/genotype). **l** Representative microscopy images of cerebral organoids stained with neuronal layer V marker CTIP2 and DAPI. The immunoreactivity of CTIP2 was quantified and normalized by DAPI fluorescent intensity (n=6 organoids/genotype). Scale bar: 50 μm. Experiments were repeated in two independently differentiated batches (i-l). All data are expressed as mean ± SEM. Student’s t tests were performed to determine statistical significance. *p<0.05, **p<0.01, ***p<0.001, **** p<0.0001. For raw data see Figure 2-source data 1 and Figure 2-source data 2.

### Activated Wnt/β-catenin signaling in astrocytes from *APOE* deficient cerebral organoids

In astrocyte clusters, “*Axonal guidance signaling*” and “*Synaptogenesis signaling*” were identified as the top ranked pathways enriched by DEGs (**Figure 2h**), indicating the important role of astrocyte-derived apoE in neuronal development and maintenance. In addition, the Wnt/β-catenin signaling pathway was also predominantly affected by *APOE* deficiency in radial glia and astrocyte populations (**Figure 2f, h**). To validate the scRNA-seq results, astrocytes were isolated from the dissociated cerebral organoids by FACS via GLAST1-mediated sorting (**Figure 3a**). Whereas *SFRP1* encoding a Wnt signaling modulator, soluble frizzled-related protein 1 (SFRP1), was one of the most significantly changed genes by *APOE* deletion in radial glia (**Figure 2e**) and astrocyte clusters (**Figure 2h**), we confirmed that *APOE* deficiency reduced *SFRP1* expression in the isolated astrocytes by RT-qPCR (**Figure 3b**). Western blotting also showed the decreased levels of SFRP1 as well as increase of β-catenin and Wnt7b in the isolated *APOE^-/-^* astrocytes compared to the controls (**Figure 3c-f**). Since SFRP1 is secreted by glia cells and has been shown to regulate neural differentiation by modulating Wnt signaling (38, 39), these results suggest that *APOE* deficiency may also impact cellular composition in the iPSC-derived cerebral organoids through Wnt/β-catenin signaling pathway by suppressing astrocytic SFRP1 production.

**Figure 3.**
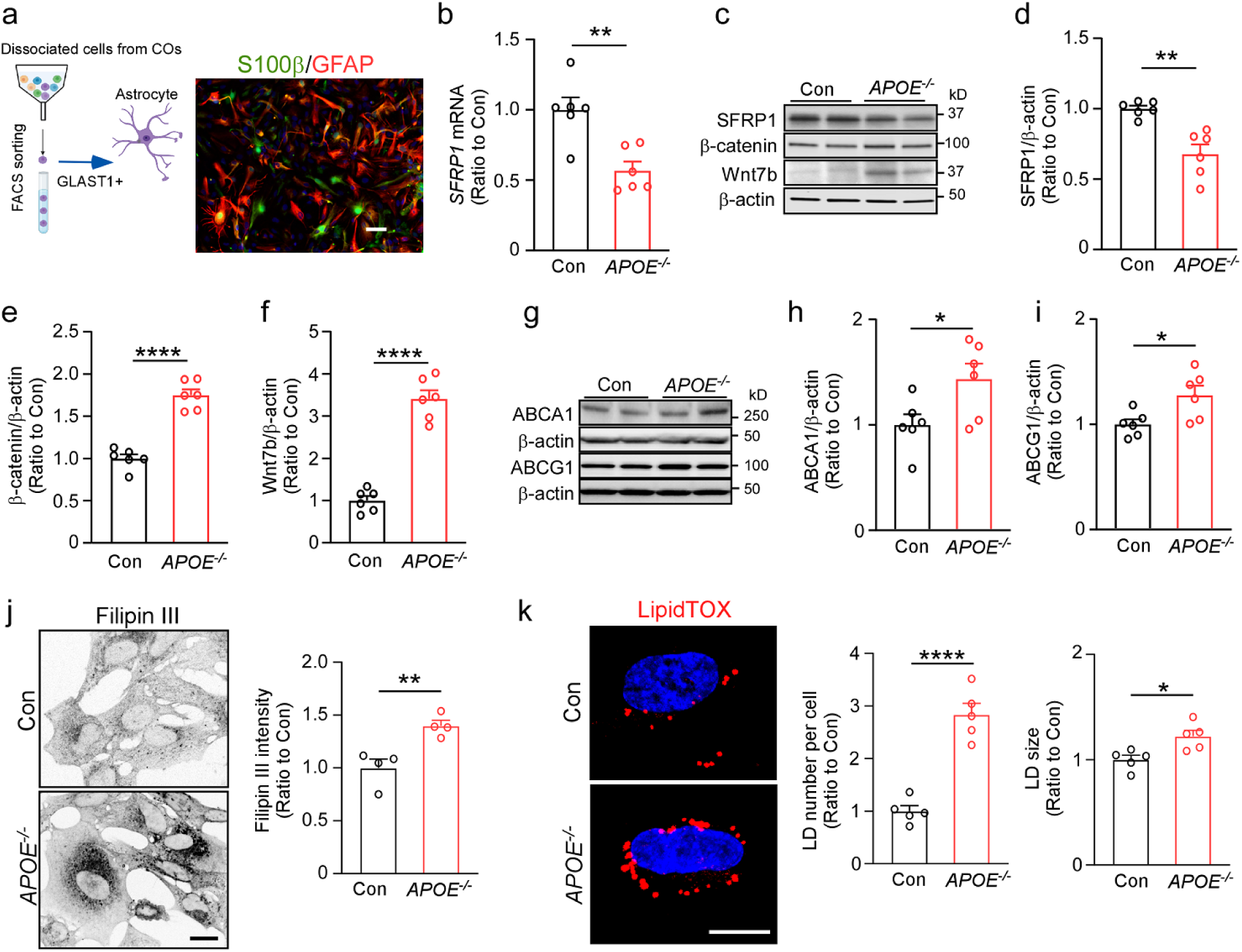
Cholesterol and lipid droplet accumulation are increased in astrocytes isolated from *APOE* deficient iPSC-derived cerebral organoids. **a** Astrocytes were isolated from the cerebral organoids through FACS using the cell surface marker GLAST1. Representative fluorescence microscopy images of the isolated astrocytes stained with antibodies against S100β and GFAP. Scale bar: 20 μm. b *SFRP1* mRNA expression in the isolated astrocytes was quantified by RT-qPCR (n=6 wells/genotype). **c-f** Protein levels of SFRP1 (d), β-catenin (e) and Wnt7b (f) in the isolated astrocytes were quantified by Western blotting (n=6 wells/genotype). **g-i** Protein levels of ABCA1 (h) and ABCG1 (i) in the isolated astrocytes were quantified by Western blotting (n = 4 wells/genotype). **j** The isolated astrocytes were plated on coverslips and stained with Filipin III for cholesterol. Filipin III intensities were quantified in 3 fields of each coverslip and averaged (n=4 coverslips/genotype), Scale bars, 20 μm. **k** The isolated astrocytes were plated on coverslips and stained with LipidTox for lipid droplets. The lipid droplet number and size per cell were quantified in 3 fields of each coverslip and averaged (n=5 coverslips/genotype). Scale bars, 10 μm. Experiments were repeated in two independently differentiated batches. All data are expressed as mean ± SEM. Student’s t tests were performed to determine statistical significance. **p<0.01, **** p<0.0001. For raw data see Figure 3-source data 1 and Figure 3-source data 2.

### Cell type-specific modification of cholesterol biosynthesis pathway in *APOE* deficient cerebral organoids

While ABCA1 and ABCG1 play important roles in lipid and cholesterol efflux (7, 40), scRNA-seq found increased expression of *ABCA1* and *ABCG1* in the astrocyte clusters of *APOE^-/-^* cerebral organoids (**Figure 2g**). Western blotting showed significant increases in the protein levels of ABCA1 and ABCG1 in astrocytes isolated from *APOE^-/-^*cerebral organoids (**Figure 3g-i**). However, greater cholesterol accumulation (**Figure 3j**) and increased lipid droplet number/size (**Figure 3k**) were observed in the *APOE^-/-^* astrocytes compared to controls when stained for Filipin III and LipidTOX, respectively. Thus, increases of *ABCA1* and *ABCG1* gene expression in *APOE* deficient astrocytes may be induced as a compensatory mechanism against excess intracellular lipid accumulation.

On the other hand, we observed an enrichment of lipid metabolism and cholesterol biosynthesis pathways in the 5 clusters of excitatory neurons, especially in cluster 3, with key cholesterol synthesis-related genes significantly upregulated (**Figure 2c-d, Supplementary Figure 4**). To validate these scRNA-seq results, excitatory neurons were isolated from the dissociated cerebral organoids by FACS via CD90-mediated sorting, where the purity of isolated neurons was confirmed by immunostaining of neuronal marker (Tuj1) (**Figure 4a**). Consistent with the results from scRNA-seq, RT-qPCR showed significant increases in the mRNA levels of multiple enzymes involved in the cholesterol biosynthesis pathway in isolated neurons from *APOE^-/-^* cerebral organoids (**Figure 4b**). Nonetheless, when intracellular cholesterol was stained with Filipin III (41) in the isolated neurons, there were no significant changes between control and *APOE^-/-^* neurons (**Figure 4c**).

**Figure 4.**
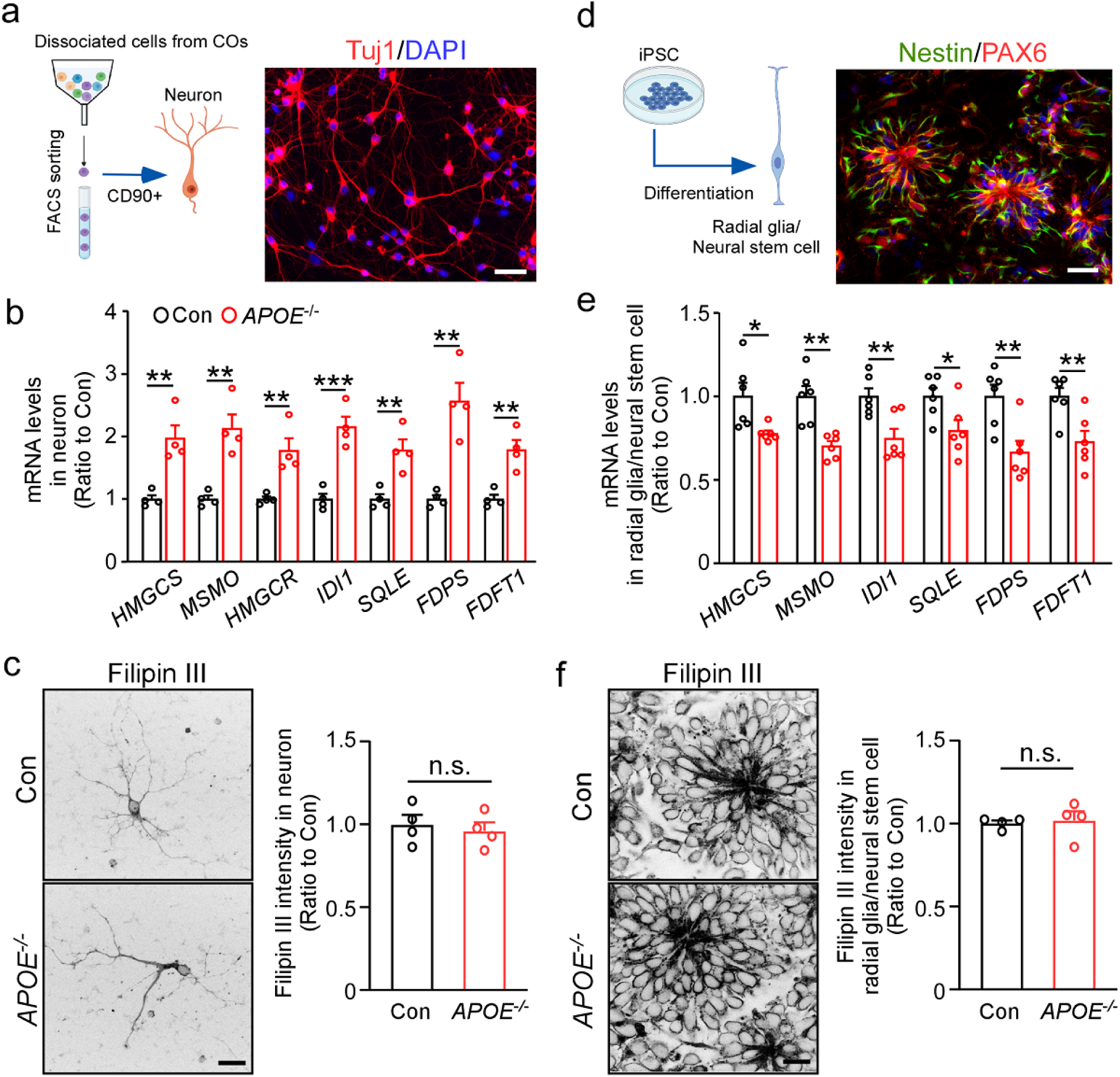
Differential effects of *APOE* deficiency on gene expression related to cholesterol biosynthesis in excitatory neurons and radial glia. **a** Neurons were isolated from the cerebral organoids through FACS using the cell surface marker CD90. Representative fluorescence microscopy images of the isolated neurons stained with antibodies against Tuj1. Scale bar: 20 μm. **b** The mRNA levels of selective cholesterol biosynthesis genes in the isolated neurons was quantified by RT-qPCR (n=4 wells/genotype). **c** The neurons isolated from cerebral organoids were plated on coverslips and stained with Filipin III for cholesterol. Filipin III intensities were quantified in 3 fields of each coverslip and averaged (n = 4 coverslips/genotype), Scale bars, 20 μm. **d** Parental and *APOE^-/-^* isogenic iPSCs were differentiated into radial glia/neural stem cells. Representative fluorescence microscopy images of the differentiated radial glia/neural stem cell stained with antibodies against Nestin and PAX6. Scale bar: 20 μm. **e** The mRNA levels of selective cholesterol biosynthesis genes in the radial glia/neural stem cell was quantified by RT-qPCR (n=6 wells/genotype). **f** The radial glia/neural stem cells were plated on coverslips and stained with Filipin III for cholesterol. Filipin III intensities were quantified in 3 fields of each coverslip and averaged (n=4 coverslips/genotype). Scale bars, 20 μm. Experiments were repeated in two independently differentiated batches. All data are expressed as mean ± SEM. Student’s t tests were performed to determine statistical significance. **p<0.01, *** p<0.001, n.s., not significant.

In contrast to the excitatory neuron population, we found that genes related to cholesterol biosynthesis were rather downregulated in both radial glia and astrocyte population (**Supplementary Figure 5**). Since radial glial cells have been identified as adult neural stem cells in the subventricular zone, which is the major source of neurons and astrocytes during development (42), we differentiated the iPSCs into radial glia/neural stem cells to validate the scRNA-seq results. Immunostaining of specific markers (Nestin and PAX6) showed the successful radial glial differentiation (**Figure 4d**). Consistent with the scRNA-seq results, mRNA levels of multiple cholesterol synthesis genes decreased in *APOE^-/-^* radial glia/neural stem cells when analyzed by RT-qPCR (**Figure 4e**), although Filipin III staining did not detect significant differences between control and *APOE^-/-^*iPSC-derived glia/neural stem cells (**Figure 4f**).

Taken together, these results suggest that *APOE* deficiency differently impacts cholesterol metabolism depending on cell types, which may influence neuronal differentiation in the iPSC-derived cerebral organoids. Repeated experiments using another set of parental and *APOE^-/-^* isogenic lines confirmed the major findings including neurogenesis deficits and lipid metabolism dysregulation due to *APOE* deficiency (**Supplementary Figure 6**).

### Altered neurogenesis and cholesterol metabolism in APOE4 cerebral organoids

*APOE4* has been associated with cell type-specific functional abnormalities in AD brain, including neurogenesis deficits, impaired synaptic function, neuronal degeneration, cholesterol lipid metabolism dysfunction and inflammatory response (9, 31, 41, 43). Thus, we explored the *APOE4* effects using the iPSC-derived cerebral organoids from an AD patient carrying *APOE ε4/ε4* (*APOE4*) and the corresponding *APOE ε3/ε3* (*APOE3*) isogenic line at Day 90. While evaluating the cellular composition differences between *APOE3* and *APOE4* organoids, RT-qPCR found decreased neuronal lineage marker mRNAs and higher astrocytic marker mRNAs in *APOE4* organoids compared to those in *APOE3* organoids (**Figure 5a**). In addition, Western blotting showed enhanced eIF2α phosphorylation levels in the *APOE4* organoids (**Figure 5b**). When neurons and astrocytes were isolated from the organoids via FACS sorting, we found that the expression of selected genes related to cholesterol biosynthesis was increased in neurons (**Figure 5c**) but decreased in astrocytes (**Figure 5d**) from *APOE4* organoids compared to those from *APOE3* organoids by RT-qPCR. Filipin III staining showed higher cellular cholesterol levels in both neurons (**Figure 5e**) and astrocytes (**Figure 5f**) isolated from *APOE4* organoids compared to those from *APOE3* organoids. Taken together, these results indicate that *APOE4* causes similar phenotypes to *APOE* deficiency in the iPSC cerebral organoids regarding neurogenesis and cholesterol metabolism, which can largely be reversed by genome editing to *APOE3*.

**Figure 5.**
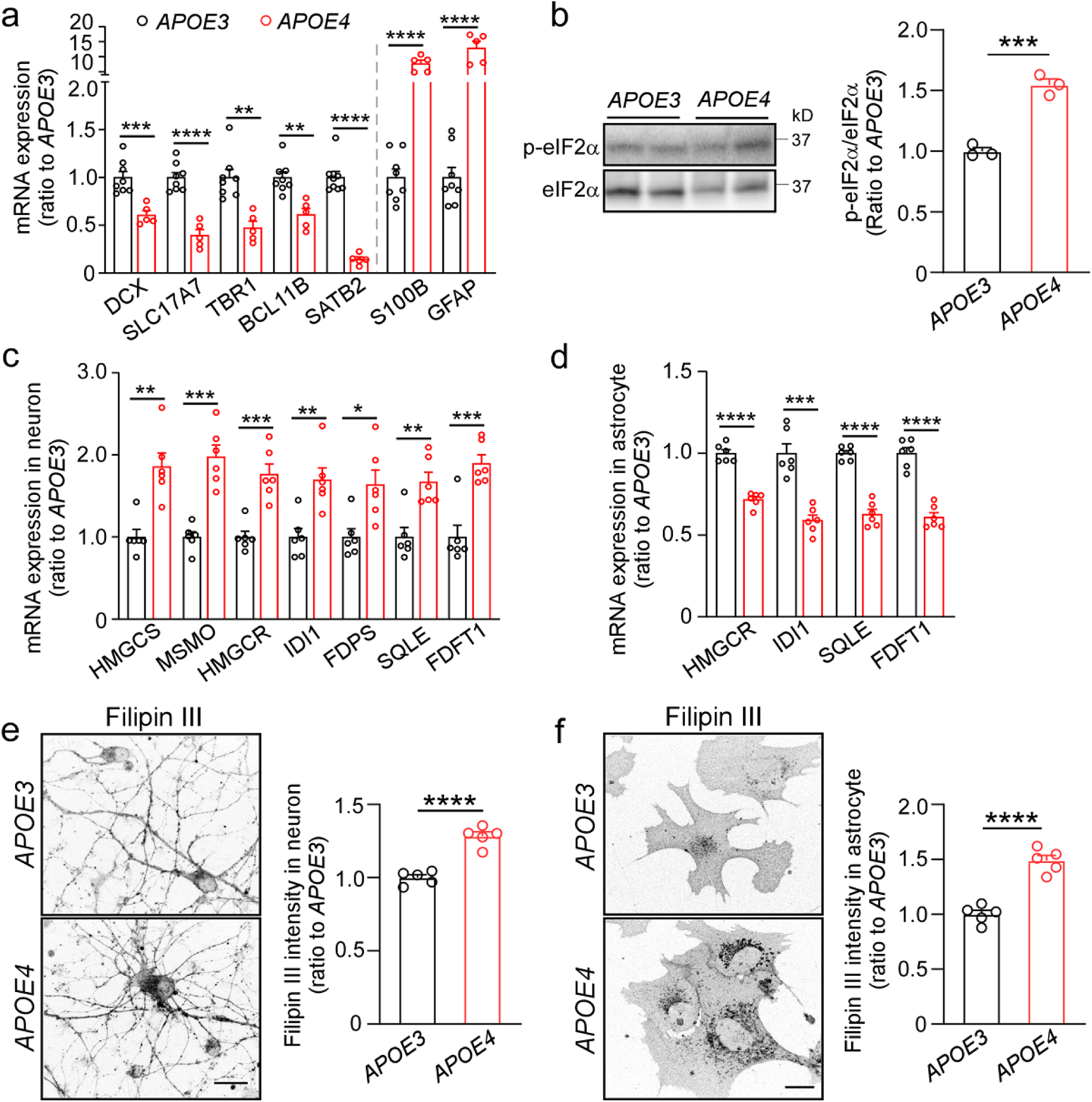
*APOE4* induces loss-of-function phenotypes in neurogenesis and cholesterol metabolism in the iPSC-derived cerebral organoids. The iPSC-derived cerebral organoids from an AD patient carrying *APOE ε4/ε4* and the *APOE ε3/ε3* isogenic line were analyzed at Day 90. **a** The mRNA levels of cerebral layer markers (*DCX*, *SLC17A7*, *TBR1*, *BCL11B* and *SATB2*) and astrocytic markers (*S100B* and *GFAP*) were quantified by RT-qPCR. Three cerebral organoids were pooled and analyzed as one sample (n=5-8 samples/genotype). **b** Phosphorylation levels of eIF2α in *APOE3* and *APOE4* cerebral organoids at Day 90 were quantified by Western blotting. Three cerebral organoids were pooled and analyzed as one sample (n=3 samples/genotype). **c, d** The mRNA levels of selective cholesterol biosynthesis genes in the neurons (c) and astrocytes (d) sorted from cerebral organoids were quantified by RT-qPCR (n=6 wells/genotype). **e, f** The isolated neurons (e) and astrocytes (f) were plated on coverslips and stained with Filipin III for cholesterol. Filipin III intensities were quantified in 5 fields of each coverslip and averaged (n=5 coverslips/genotype), Scale bars, 20 μm. Experiments were repeated in two independently differentiated batches. All data are expressed as mean ± SEM Student’s t tests were performed to determine statistical significance. *p<0.05, **p<0.01, ***p<0.001, **** p<0.0001. For raw data see Figure 5-source data 1.

## Discussion

The iPSC-derived cerebral organoids provide a unique model system in which human cell type specification, self-organization and heterogeneous intercellular communication occur simultaneously. In this study, we investigated how *APOE* deficiency influences the molecular pathways and cellular functions in the organoids by scRNA-seq combined with multiple validation approaches. While several cell types were identified in cerebral organoids consistent with previous findings (29, 34), we found that *APOE* deficiency dramatically modulates the cellular compositions, accompanied by enhanced cellular stress and lipid dyshomeostasis. Our scRNA-seq data revealed an increase of radial glia and astrocyte populations, but a decrease of excitatory neurons in isogenic *APOE^-/-^* cerebral organoids. Furthermore, we found a significant increase of inhibitory neurons in *APOE^-/-^* cerebral organoids. Interesting, such developmental excitation-inhibition imbalance has been observed in several other disease-related iPSC cerebral organoid model systems (44–46). Indeed, apoE has been identified as an important regulator balancing brain progenitor cell fate and regulating neurogenesis in the dentate gyrus (47–49). At the early developmental stage, apoE is essential for inhibiting cell proliferation, maintaining neural precursor characteristics and promoting neurogenesis (47). *Apoe* knockout mice also show the excessive proliferation of neural stem cells and a shift of neural differentiation from neurogenesis to astrogenesis (47, 49). Thus, it would be interesting to further define the molecular mechanisms by which apoE regulates neuronal fate, providing insight into the roles of apoE isoforms in AD and other neurological diseases.

Although the role of apoE in lipid metabolism has been intensively investigated, there is still a lack of comprehensive assessment on how apoE regulates lipid homeostasis in a cell type-specific manner. As apoE predominantly mediates lipid efflux, our results showed that *APOE* deficiency results in excess cholesterol accumulation and lipid droplet formation in astrocytes isolated form iPSC-derived cerebral organoids. The increased levels of ABCA1 and ABCG1 in the *APOE^-/-^* astrocytes are likely due to a compensative reaction to excessive intracellular lipid accumulation. Intriguingly, cholesterol has been shown to activate canonical Wnt/β-catenin signaling (50), consistent with our finding in astrocyte population through scRNA-seq. Thus, the increase of astrocyte proportion in *APOE^-/-^* cerebral organoids may be induced at least partially by the activated Wnt/β-catenin pathway in astrocytes or radial glia as Wnt/β-catenin signaling has been shown to be involved in the differentiation process of reactive astrocytes upon brain injury (51). In addition, the lipid overload status is presumably induced in radial glia/neural stem cells when apoE-mediated lipid efflux is suppressed, since the reduction of cholesterol synthesis-related genes were observed in radial glia as well as astrocyte populations in the *APOE^-/-^* organoids. While lipid droplets are highly abundant in neural stem cells under physiological conditions (52) which correlates with increased proliferative ability (53), the excess lipid accumulation in neural stem cells may steer their differentiation fate into astrocytic lineage although further studies are needed.

Since neurons rely on astrocytic apoE for cholesterol supplies to maintain their functions and homeostasis (5, 54), neuronal populations may be vulnerable to the relatively stressful condition of the organoid models in the absence of apoE. Indeed, *APOE^-/-^* cerebral organoids showed upregulation of cellular stress related pathways, including EIF2 signaling and mTOR signaling. We also found significant increase of cholesterol biosynthesis genes in excitatory neuron population in *APOE^-/-^* cerebral organoids, which likely reflects a compensatory reaction to the insufficient lipid supplies due to the lack of astrocytic apoE. Consistent with our finding, astrocytic apoE has been shown to specifically silence genes involved in neuronal cholesterol biosynthesis (5). As cholesterol biosynthesis requires multiple enzymatic steps that involve high energy consumption, the emergent condition may also exacerbate ER stress and autophagy in neurons thereby influencing the cellular composition in *APOE^-/-^*cerebral organoids. Of note, we demonstrated that the administration with eIF2 phosphorylation inhibitor ISRIB restores the altered cellular composition in the *APOE^-/-^* cerebral organoids by facilitating neurogenesis and suppressing astrogliosis. Since eIF2α phosphorylation is the central event in the integrated stress response (ISR) (55), our findings indicate the predominant involvement of ISR cause by *APOE* deficiency in defining neuronal differentiation.

ApoE4 has been shown to mediate cholesterol efflux in astrocytes less efficiently than apoE3, resulting in the exacerbated accumulation of cholesterol and lipid droplets (22, 41, 56). We also found that the expression of cholesterol synthesis gene is lower in neurons and higher in astrocytes isolated from *APOE4* cerebral organoids than those with *APOE3*, which is accompanied by the increase of intracellular cholesterol accumulation. Since *APOE4* organoids showed similar phenotypes with *APOE^-/-^*cerebral organoids, our results suggest that *APOE4* may cause lipid dyshomeostasis through loss-of-function effects compared to *APOE3*. In addition, *APOE4* cerebral organoids also possessed diminished neuronal differentiation similar to the *APOE^-/-^*organoids. Consistently, studies in mouse models have found that both *Apoe* deficiency and human apoE4 isoform impair the hippocampal neurogenesis (48, 49). Importantly, since the significant increase of the phosphorylation level of eIF2α was observed in *APOE4* cerebral organoids, our results suggest that ISR inhibition can be a potential therapeutic target for *APOE4*-mediated pathogenesis in AD. Indeed, several recent studies show that ISRIB could restore hippocampal protein synthesis and delay cognitive decline in aged mice and AD mouse models (57, 58).

In summary, our study demonstrates the essential role of apoE in brain homeostasis using the iPSC-derived cerebral organoids and offers critical insights into the underlying mechanisms of *APOE4*-related phenotypes. *APOE* deficiency and *APOE4* influence lipid metabolism and neuronal differentiation by activating ISR pathway in a cell type-specific manner. One limitation of our study is the *ex vivo* nature of iPSC-derived cerebral organoids lacking systemic effects compared to *in vivo* environment. Another limitation of the cerebral organoid system is the lack of several important brain cell types, such as microglia and vascular cells (29, 32). Thus, future studies should be directed to validating our findings in human brains and optimized iPSC-derived cerebral organoid systems with the incorporation of additional brain cell types. Together, our findings will provide novel insights on the underlying mechanisms in *APOE4*-associated disease pathogenesis and guide the development of apoE-targeted therapy.

## Material and Method

### APOE-deficient and genotype-specific isogenic iPSC lines

Two different sets of human parental iPSC lines and their isogenic *APOE^-/-^* iPSC lines were used in the study. One set of the parental (Con) and isogenic *APOE^-/-^* iPSC lines were obtained from the Xcell Science. Detailed information about these lines can be found in the Xcell Science website (http://www.xcellscience.com/products/ipsc). For the MC0192 iPSC line set, isogenic *APOE^-/-^* iPSC line was obtained via CRISPR/Cas9 knockout of the *APOE* gene. *APOE* deletion for both isogenic sets have been confirmed in our previous publication (31). The *APOE4* and *APOE3* isogenic lines were kind gifts from Dr. Yadong Huang (15).

### Cerebral organoid culture

Cerebral organoids were generated using STEMdiff™ Cerebral Organoid Kit (Stemcell Technologies) following manufacturer’s instructions. Human iPSC colonies were dissociated into single cell suspension with Accutase at Day 0. Cells were seeded into a U-bottom ultra-low-attachment 96-well plate (15,000 cells/well) in Medium A with 10 μM Y-27632. Additional 100 μL of medium A were added into each well on Day 2 and Day 4, respectively. EBs were moved to 48-well low attachment plates in Medium B on Day 5 and left for an additional 3-5 days. EBs were further embedded into 20 μL of matrigel and cultured in Medium C+D in 6-well low attachment plates for 3 days. In the final stage, organoids were moved to an orbital shaker in 10 cm dishes and cultured in Medium E, which was replaced with neuronal maturation medium after 4 weeks. Cerebral organoids were harvested at Day 90 for scRNA-seq and validation. For ISRIB treatment, cerebral organoids at Day 60 were treated with neuronal maturation medium containing 100 nM ISRIB or vesicle DMSO for 30 days. Medium was changed every three days and cerebral organoids at Day 90 were harvested for further analysis. Cerebral organoids in abnormal size (<1mm diameter) were considered unhealthy and excluded from further experiments.

### Single cell suspension, library preparation and sequencing

The parental and isogenic *APOE-/-* iPSC lines from the Xcell Science were differentiated into cerebral organoids and subjected to single cell RNA-sequencing (scRNA-seq) analysis at Day 90. Cerebral organoids were dissociated into single cell suspension using the Worthington Papain Dissociation System kit (Worthington Biochemical, LK003150) following manufacturer’s instruction (32). Cerebral organoids were gently transferred to an individual 60 mm dish with 2.5 mL of Papain + DNase solution. Three organoids were pooled as one sample. Cerebral organoids were minced into small pieces (< 1 mm) and the plates were transferred to an orbital shaker at 45 rpm inside a humidified tissue culture incubator at 37°C and 5% CO2. After 15 minutes, minced pieces were gently dissociated using a 1 mL pipette and returned to the incubator for additional 15 minutes. Dissociated tissues were gently pipetted up and down 10 times and transferred to an empty 15 mL conical tube for the debris to settle (1-3 minutes). Cell suspension (avoid debris) was transferred to the prepared stop solution and centrifuged at 300 g for 5 minutes. Dissociated cells were resuspended in ice-cold PBS containing 0.05% BSA and passed through 30 μm cell strainer. In all samples, more than 90% of the cells were viable based on trypan blue examination. Approximately 6,000 isolated cells from one sample were loaded into each sample well on a chip and combined with Gel Beads containing barcoded oligonucleotides using a 10x Chromium Controller (10x Genomics). Single cell libraries were constructed according to the manufacturer’s instructions and were sequenced by an Illumina HiSeq 4000 Sequencing Systems.

### Single-Cell transcriptome analysis

The single cell sequence preprocessing was performed using the standard 10x Genomics Cell Ranger Single Cell Software Suite (v5.0.0). Briefly, raw sequencing data were demultiplexed, aligned to the Human reference genome, GRCh38 2020-A (GENCODE v32/Ensembl 98), and the reads aligned to each gene were counted. Cell filtration, normalization, clustering, and differential expression analyses were performed with R (v4.0.3) and the single-cell analysis package Seurat v4 (59). For quality control, cells with greater than 25000 unique molecular identities (UMI) counts, less than 5000 counts, less than 200 features, and mitochondria percentage greater than 12% were discarded. The resulting count matrix was 36601 genes by 27199 cells. For each line (parental or isogenic), SCTransform was applied setting vars.to.regress to mitochondria percentage (60). Each organoid was treated as a separate batch and integrated with IntegrateData, setting normalization.method to ‘SCT’, and setting references to organoid 1 from each line. Following batch correction, FindNeighbors was run on the first 20 Principal Components (PCs), FindClusters was run with resolution = 0.5. Clusters were visualized with t-SNE plot run on the first 30 PCs. For computing marker genes of each cluster, FindMarkers was run using the MAST method, and setting latent.vars to the cellular detection rate, computed by scaling the number of features per cell (61). For computing differentially expressed genes between conditions, cells were subset either to each cell type or cell cluster, and DEGs were computed again with FindMarkers using MAST method, setting latent.vars to the cellular detection rate. DEGs were filtered for Bonferroni-corrected p values < 0.05 and absolute value of log2 fold change greater than 0.25. Pathway analyses were performed with Ingenuity Pathway Analysis (Content version: 62089861). For pseudotime trajectory analysis, Monocle3 was run with default settings, using the previously computed t-SNE plot dimensionality reduction as input (62). For computing the trajectory graph, the root node was manually selected to correspond to cells within cluster 4 radial glia. Cell types that were not part of the trajectory were removed for visualization.

### Neuron and astrocyte isolation by FACS

Cerebral organoids were dissociated into single cells following the same cell isolation protocol described above. All steps were performed using sterile techniques. Dissociated cells were resuspended in ice-cold PBS containing 0.05% BSA and passed through 30 μm cell strainer. Cell suspension was then centrifuged at 300 g for 5 minutes at 4°C. Cell pellet was resuspended in basal medium plus 0.5% BSA (basal medium: phenol-red-free DMEM/F12 + Neurobasal + 1 x N2 + 1 x B27) and incubated on ice for 15 minutes. Cells were washed with basal medium and resuspended at a concentration of 1 × 10^7^ cells/mL in basal medium plus 0.05% BSA in a 5 mL round bottom tube. Human anti-CD90-PE antibody (Miltenyi Biotec, 130-117-388) and anti-GLAST-APC antibody (Miltenyi Biotec, 130-123-641) were added according to manufacturers’ recommendations and incubated on ice for 30 minutes. Stained cells were washed with basal medium for three times, centrifuged at 300 g for 5 minutes, and resuspended with basal medium plus 0.05% BSA at 2.5x10^6^ cells/ml for FACS sorting (BD FACS Aria). Sytox Blue (1:1000 dilution, ThermoFisher S34857) was used as the viability dye. Cells were sorted on a FACS Aria II (BD Biosciences) equipped with 405 nm, 561 nm, and 633 nm lasers, using the 100 micron nozzle with 20 psi sort pressure. In preliminary and antibody titration experiments, FMO and single-stain controls were found to be equivalent, therefore sort gates were drawn based on single-stained controls. Sort targets were determined by identifying cells on FSC-A vs SSC-A, then performing doublet discrimination with hierarchical SSC-H vs. SSC-W and FSC-H vs. FSC-W plots, then identifying lives cells on a Sytox Blue-BV421-A vs. FSC-A plot, and finally interrogating GLAST-APC-A vs. CD90-PE-A for CD90 single positive and GLAST single positive cells. CD90+ neurons were harvested for RT-QPCR analysis or seeded on PLO/Laminin coated coverslips for downstream assays. GLAST+ astrocytes were plated in Matrigel coated wells in astrocyte media (ScienCell) and expanded for downstream assays. The purity of sorted cells was confirmed by immunostaining of cell-type-specific markers.

### Radial glia/Neural stem cell differentiation

Radial glia/neural stem cell were generated using STEMdiff™ SMADi Neural Induction Kit (Stemcell Technologies, 08581) following manufacturer’s instruction. Human iPSCs were dissociated into single cells and seeded on AggreWell 800 plates in EB formation medium (Stemcell Technologies, 05893) to initiate EB formation. After 24 hours, EB formation medium was exchanged to Neural induction medium with daily half medium change for 3-4 days. Next, EBs were collected and replated onto Matrigel-coated dishes and cultured in neural induction medium for another 5-7 days to induce neural rosette formation. Neural rosettes were isolated as a single cell suspension and re-plated onto Matrigel-coated dishes in neural induction medium for another 2-3 days. Radial glia/Neural stem cells were maintained and amplified in Neural progenitor cell medium (Stemcell Technologies, 05833) for further experiments.

### Tissue processing

Cerebral organoids or cells were lysed with RIPA Lysis and Extraction Buffer supplemented with Protease and Phosphatase Inhibitor Cocktails for Cell Lysis (Roche). Samples were kept on ice for 60 minutes after sonication, and then centrifuged in an ultracentrifuge (Beckman-Coulter) at 100,000 g, for 1 hour at 4°C. Supernatants were collected for Western blotting analysis. Total protein concentration in the soluble fraction was determined using a Pierce BCA Protein Assay Kit.

### Immunofluorescence staining

Cerebral organoids and specific cell types were harvested and fixed in 4% paraformaldehyde for 30 minutes then washed with PBS three times. After fixation, cerebral organoids were dehydrated with 30% sucrose in PBS at 4°C. Optical cutting temperature (OCT) compound (VWR) was used to embed cerebral organoids and frozen on dry ice. Tissue was sectioned at 30 μm and collected on glass slides and stored at -20°C. For immunostaining, tissue sections or fixed cells were permeabilized in 0.25% Triton X-100 and blocked with blocking buffer containing 4% normal donkey serum, 2% BSA and 1 M glycine in PBS. Sections were then incubated with primary antibodies in blocking buffer overnight at 4°C. Primary antibodies and their dilutions used in this study are as follows: Tuj1 (Sigma, T2200, 1:1000), GFAP (Millipore, MAB360, 1:300), S100β (Abcam, ab52642, 1:100), Nestin (Abcam, ab18102, 1:500), Pax6 (Biolegend, 901302, 1:300) and CTIP2 (Abcam, ab18465, 1:100). On the following day, sections were washed three times with PBS, and then incubated with secondary antibodies for 2 hours at room temperature. Finally, sections were washed three times with PBS before mounting with the glass coverslip. After washing three times with PBS, samples were incubated with fluorescently conjugated secondary antibodies (Alexa Fluor 488 and 594 conjugates, Invitrogen, 1:500) for 2 hours at room temperature and washed three times with PBS before mounting with the glass coverslip. Sections were washed three times in PBS and mounted with Vectashield (H-1000, Vector Laboratories). Fluorescent signals were detected by Keyence fluorescence microscopy (model BZ-X, Keyence) and quantified using ImageJ software.

### Filipin III and LipidTOX staining

Cholesterol Assay Kit (Cell-Based) (Abcam, ab133116) and HCS LipidTOX™ Red Neutral Lipid Stain (Invitrogen, H34476) were used for cholesterol and lipid droplet detection, respectively. Cells were fixed using 4% PFA for 30 minutes, then incubated with Filipin III or LipidTOX solution prepared according to manufacturer’s recommendations for 60 minutes and examined immediately on a confocal laser scanning fluorescent microscopy (model LSM880 Invert, Carl Zeiss) using 40× oil (EC Plan-Neofluar 40x/1.30 Oil DIC M27) or 63× oil (Plan-Apochromat 63x/1.40 Oil DIC M27) objectives with numerical aperture of 1.0. For visualization, excitation of Filipin was at 405 nm and LipidTOX was excited at 594 nm. The Filipin average intensities were quantified using Fiji software. Lipid droplets were analyzed with custom scripts written in Matlab (r2020b). Briefly, images were thresholded with constant thresholds for each individual channel. Droplets were detected by binarizing images, applying area filter, applying Matlab’s imfindcircles function, then applying watershed transform on selected circular features.

### Western blotting

RIPA fractions collected from cerebral organoids were run on a 4-20% sodium dodecyl sulfate-polyacrylamide gel (Bio-Rad), and transferred to PVDF Immobilon FL membranes (Millipore). After blocking in 5% non-fat milk in PBS, membranes were incubated overnight with primary antibodies in 5% non-fat milk/PBS containing 0.1% Tween-20. Primary antibodies and their dilutions used in this study are as follows: ABCA1 (Millipore, MAB10005, 1:1000), ABCG1 (Abcam, ab52617, 1: 1000), PhosphoPlus eIF2α (Ser51) Antibody Duet (Cell Signaling Technology, 89117, 1:1000), Sfrp1 (Invitrogen, MA5-38193, 1:1000), β-catenin (BD Biosciences, 610154, 1:4000), Wnt7B (Abcam, ab227607, 1:1000) and β-actin (Sigma, A2228, 1:4000). After 24 hours, membranes were probed with LI-COR IRDye secondary antibodies or horseradish peroxidase-conjugated secondary antibody for 2 hours, which was further detected with SuperSignal West Femto Chemiluminescent Substrate (Pierce).

### RT-qPCR

Trizol/chloroform method was used to extract RNA from organoids and different cell types, followed by DNase and cleanup using the RNase-Free DNase Set and the RNeasy Mini Kit (QIAGEN). The cDNA was prepared with the iScript cDNA synthesis kit (Bio-Rad). Real-time qPCR was conducted with Universal SYBR Green Supermix (Bio-Rad) using an iCycler thermocycler (Bio-Rad). Relative gene expression was normalized to *ACTB* gene coding β-actin and assessed using the 2−ΔΔCT method. Primers used to amplify target genes by RT-qPCR are as **Supplementary Table 1**.

### SFRP1 ELISA

The SFRP1 level in the medium was measured using the Human SFRP1 ELISA Kit (Abcam, ab277082) according to the manufacturer’s instruction. Diluted samples or standards (50 µL/well) were added to the plates, followed by 50 µL of the Antibody Cocktail to each well. The plates were sealed and incubated for 1 hour at room temperature on a plate shaker set to 400 rpm. The plates were washed three times and incubated with 100 µL of TMB Development Solution to each well for 10 minutes. The reaction was stopped and read at 450 nm with a microplate reader (Biotek). Results were normalized to total protein concentration of the cell lysate.

### Statistical Analyses

For cerebral organoid comparison between two groups, the student’s t test was performed to determine the significance using GraphPad Prism. All statistical tests were two-sided. Data were presented as Mean ± SEM. A p value of < 0.05 was considered statistically significant. Specific statistical methods, the number of the experiments, and the significance levels for each analysis are described in the legends of individual figures.

## Acknowledgement

This work was supported by NIH grants U19AG069701, RF1AG057181, R37AG027924, RF1AG046205, P01NS074969, and U54NS110435 (to G.B.), a Cure Alzheimer’s Fund grant (to G.B.), an Alzheimer’s Association Research Fellowship 2018-AARF-592302 (to J.Z.), and NIH grant R01AG061796 (to N.E.-T.). This work was also partially supported by Mayo Clinic Center for Regenerative Medicine, Neuroregeneration Lab. We are grateful to Dr. Yadong Huang and Dr. Chengzhong Wang for providing the *APOE4* and isogenic *APOE3* iPSC lines.

## Author contributions

J.Z., N.E.-T., T.K. and G.B. conceived and designed the project, and wrote the paper. Y.M. helped with collecting human skin biopsies and generating iPSC lines. J.Z., W.L., J.M., Y.L., L.L. T.I. and Y.Z., executed the experiments and analyzed the data. T.I., Y.R. and Y.A. performed analysis for RNA-Sequencing data. All authors reviewed and approved the final draft of the manuscript.

## Competing interests

G.B. consults for SciNeuro, has consulted for AbbVie, E-Scape, Eisai, and Vida Ventures, is on the scientific advisory board for Kisbee Therapeutics. All other authors declare no competing interests.

## Data availability

The single cell RNA-seq data are available via the AD Knowledge Portal (https://adknowledgeportal.synapse.org, Data reference number: syn30866487). The AD Knowledge Portal is a platform for accessing data, analyses, and tools generated by the Accelerating Medicines Partnership (AMPAD) Target Discovery Program and other National Institute on Aging (NIA)-supported programs to enable open-science practices and accelerate translational learning. The data, analyses, and tools are shared early in the research cycle without a publication embargo on secondary use. Data are available for general research use according to the following requirements for data access and data attribution [https://adknowledgeportal.synapse.org/ DataAccess /Instructions]. All other data that support the findings of this study are available from the corresponding authors upon reasonable request.

## Supplementary information

**Supplementary Figure 1.**
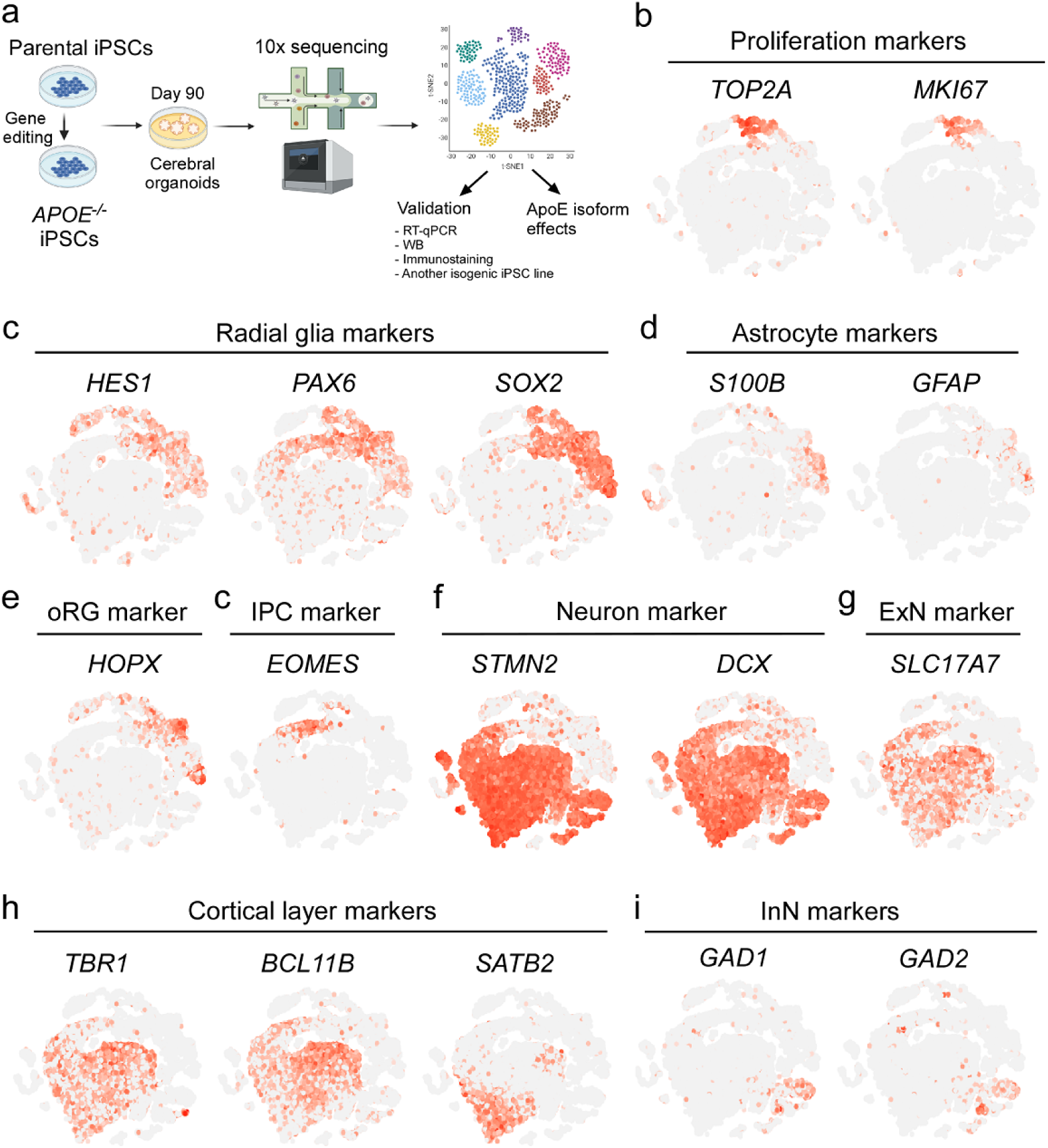
Marker gene expression in iPSC-derived cerebral organoids. **a** The schematic shows the workflow of scRNA-seq analysis of cerebral organoids and related validation. **b-i** t-SNE plots indicating the expression of key marker genes for different cell types. oRG, outer radial glia; IPC, intermediate progenitor cell; ExN, excitatory neuron; InN, inhibitory neuron.

**Supplementary Figure 2.**
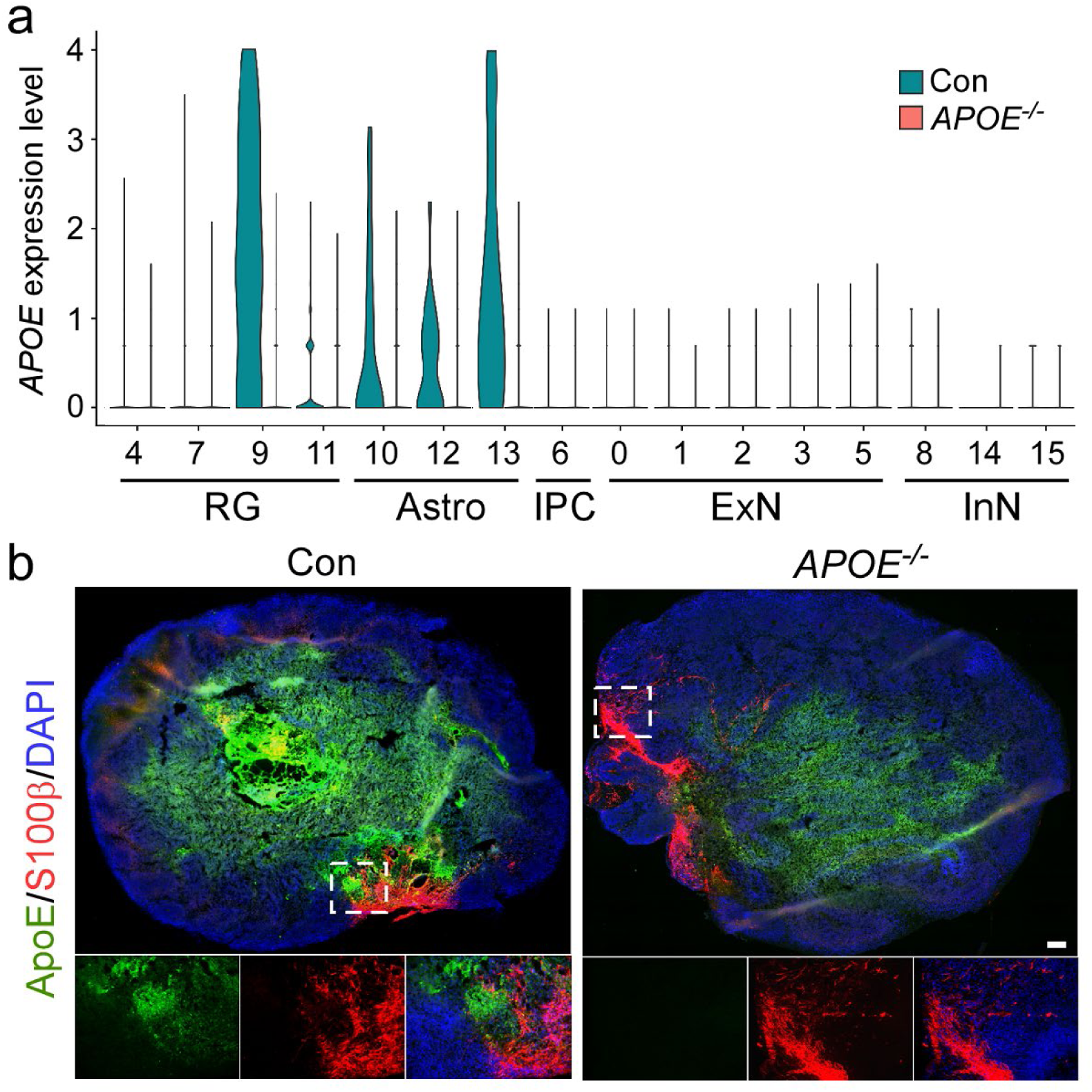
*APOE* expression and distribution in the iPSC-derived cerebral organoids. **a** Violin plot of *APOE* expression in different cell clusters of both control and *APOE^-/-^* cerebral organoids. **b** Representative images of immunostaining for apoE and an astrocytic marker S100β in cerebral organoids. Scale bar: 200 μm.

**Supplementary Figure 3.**
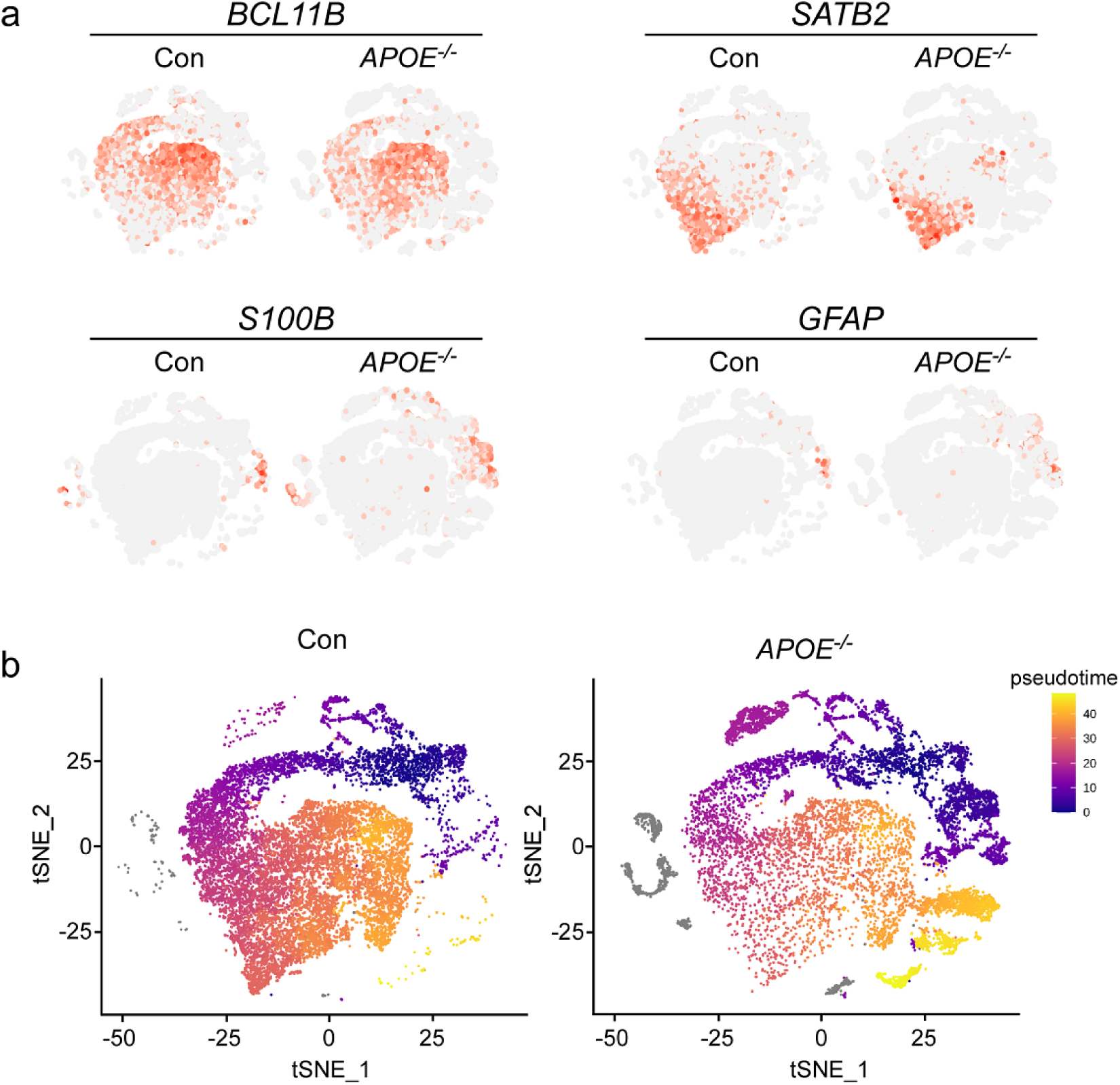
Differentiation pattern changes in the *APOE* deficient cerebral organoids. **a** t-SNE plots for cerebral layer markers (*BCL11B*, *SATB2*) and astrocytic markers (*S100B*, *GFAP*) in the control and *APOE^-/-^*cerebral organoids. **b** Pseudotime trajectory analysis of the cerebral organoids. Cells (dots) are colored according to pseudotime, from origin in dark purple to terminal state in light yellow.

**Supplementary figure 4.**
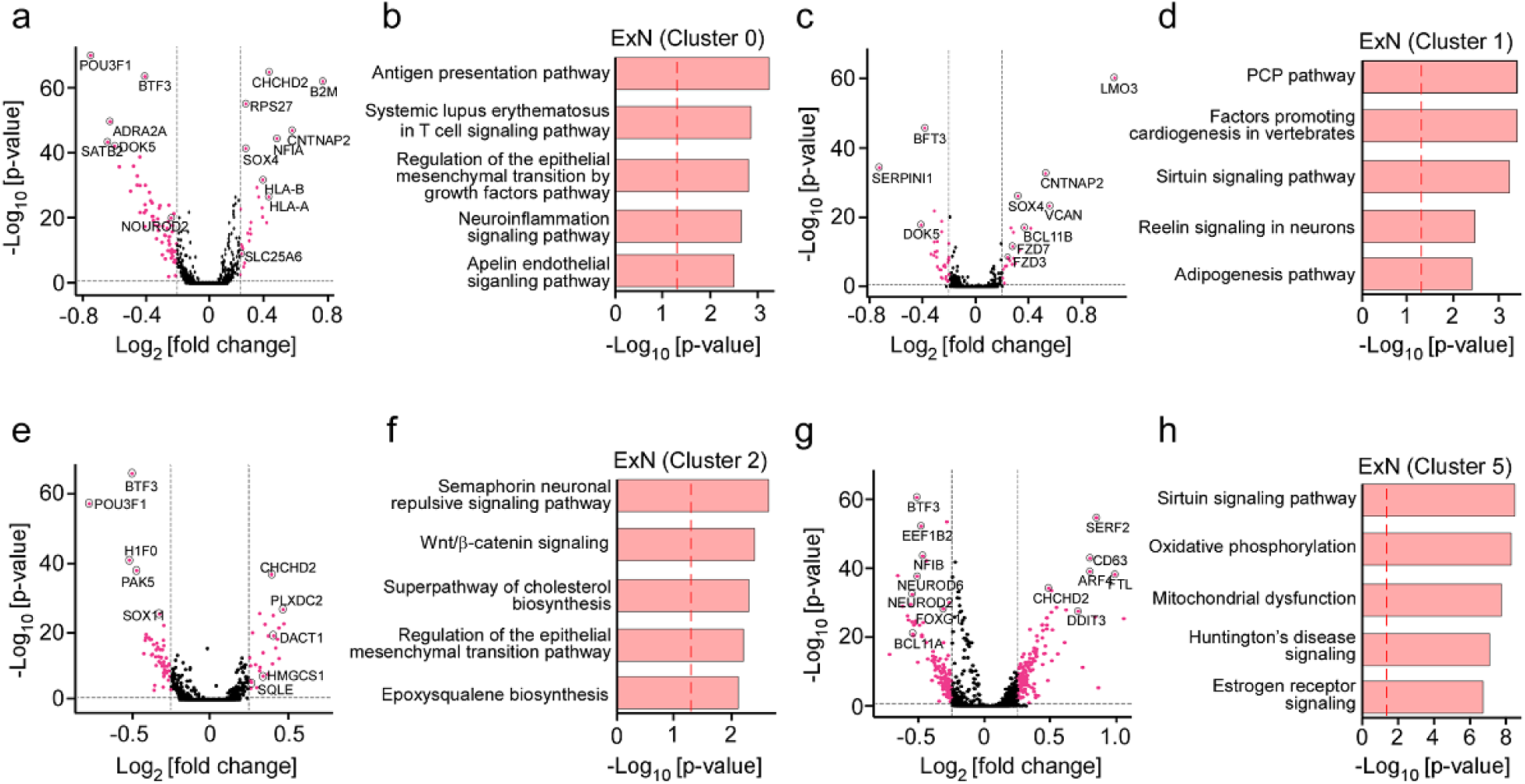
DEGs and pathway analysis for each excitatory neuron cluster. Volcano plots for DEGs and gene ontology analyses in excitatory neuron cluster 0 (a, b), cluster 1 (c, d), cluster 2 (e, f), and cluster 5 (g, h).

**Supplementary Figure 5.**
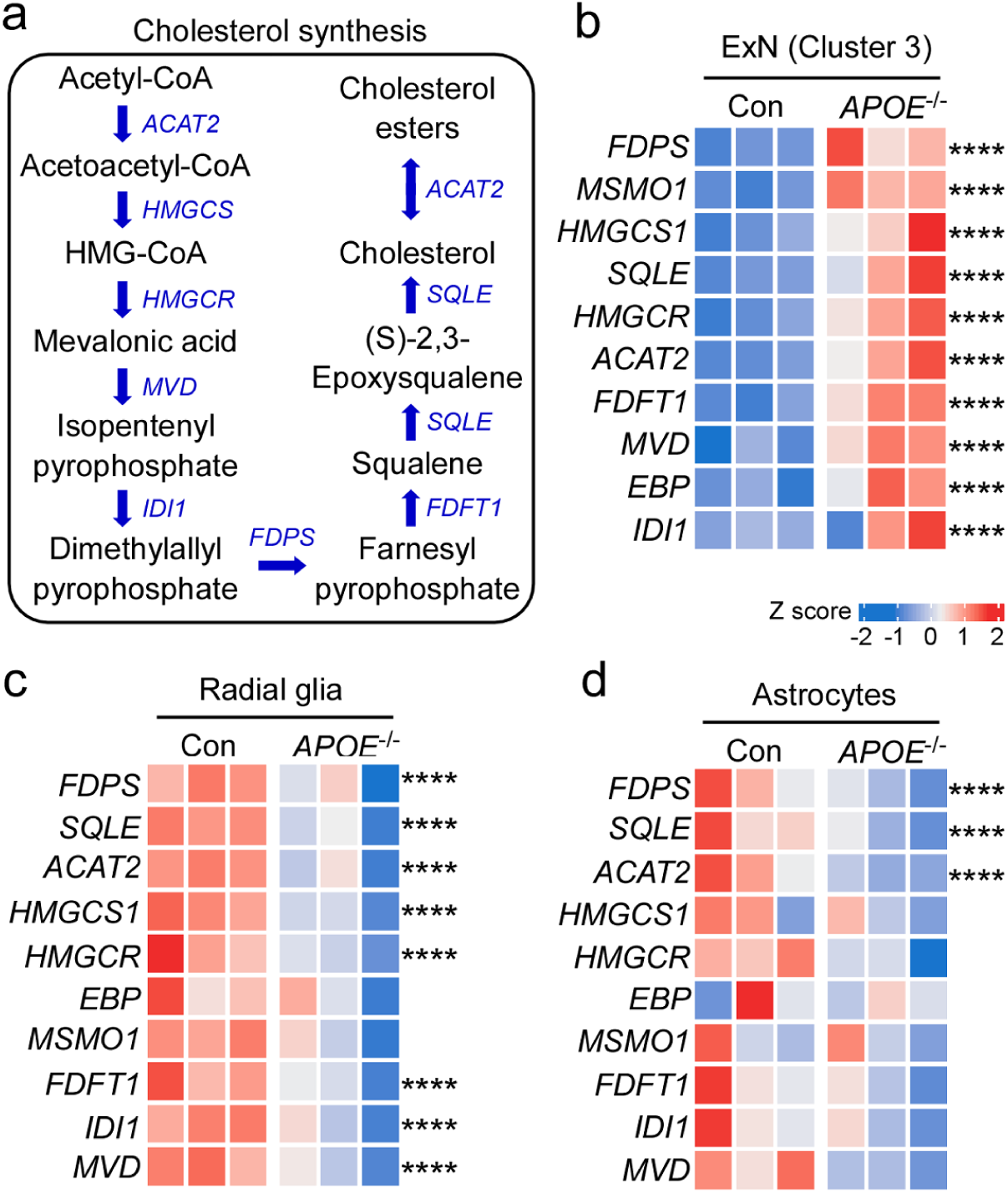
Changes in the expression of cholesterol biosynthesis-related genes in different cell types within the iPSC-derived cerebral organoids. **a** Schematic diagram for cholesterol biosynthesis in Bloch pathway. **b-d** Expression of specific cholesterol biosynthetic genes in excitatory neurons cluster 3 (b), radial glia (c) and astrocytes (d) from the control and *APOE^-/-^* cerebral organoids are visualized in a heatmap.

**Supplementary Figure 6.**
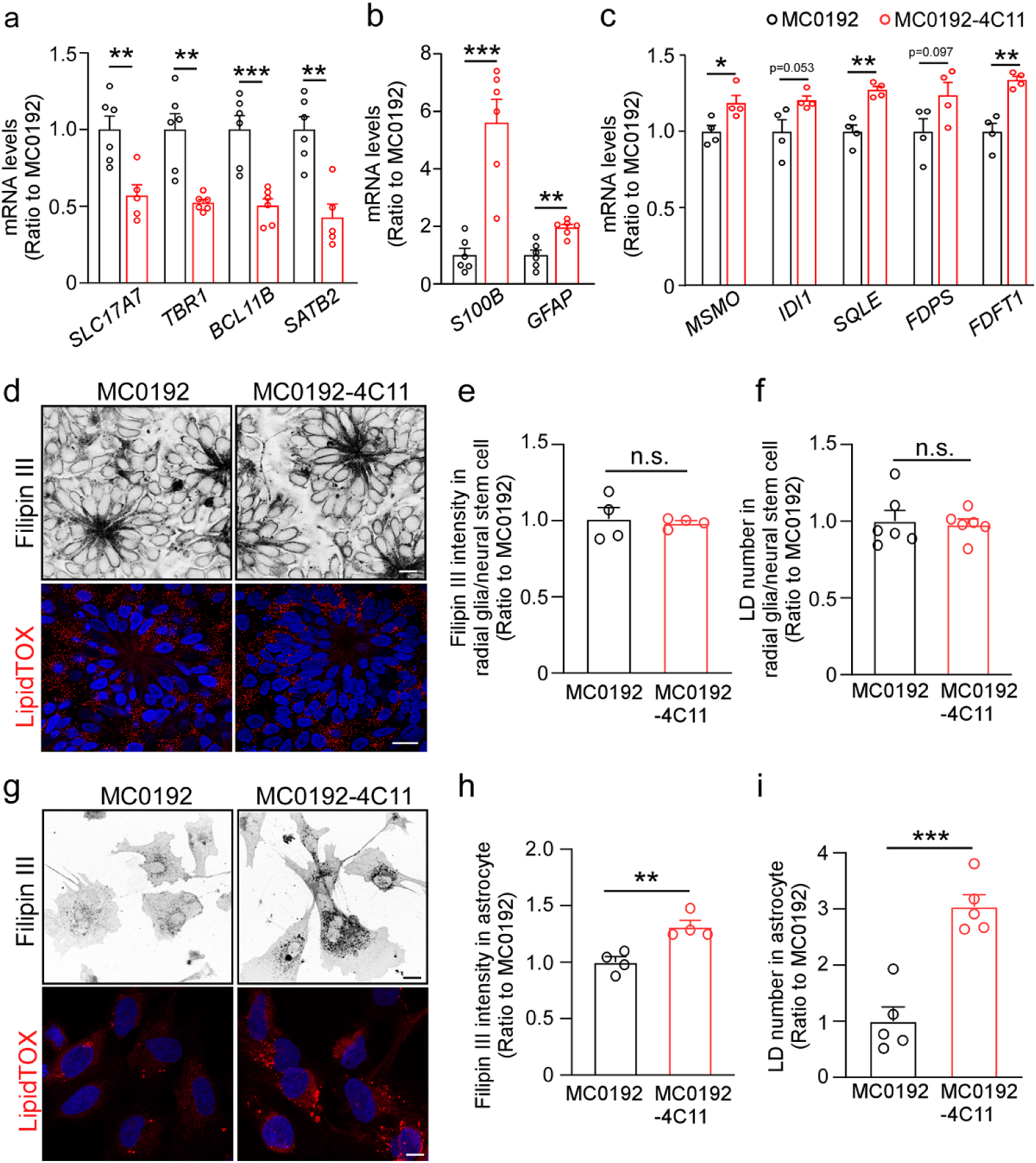
Confirmation of key findings using another set of *APOE* deficient iPSC-derived cerebral organoids. Another *APOE* deficient iPSC line (MC0192-4C11) was generated from a control iPSC line (MC0192). The parent control and isogenic *APOE* deficient iPSCs were differentiated into cerebral organoids and subjected to analysis at Day 90. **a, b** The mRNA levels of cerebral layer markers (a; *SLC17A7*, *TBR1*, *BCL11B* and *SATB2*) and astrocytic markers (b; *S100B* and *GFAP*) were quantified by RT-qPCR. Three cerebral organoids were pooled and analyzed as one sample (n=6 samples/genotype). **c** The mRNA levels of selective cholesterol biosynthesis genes in neurons isolated from the cerebral organoids were quantified by RT-qPCR (n=4 wells/genotype). **d-f** The radial glia/neural stem cells differentiated from the iPSCs were plated on coverslips and stained with Filipin III and LipidTOX (d). Filipin III intensities (e) and lipid droplet number (f) were quantified in 3 fields of each coverslip and averaged (n=4-6 coverslips/genotype). Scale bars, 10 μm. **g-i** The isolated astrocytes from the iPSCs were plated on coverslips and stained with Filipin III and LipidTOX (g). Filipin III intensities (h) and lipid droplet number (i) were quantified in 3 fields of each coverslip and averaged (n=4-5 coverslips/genotype). Scale bars, 10 μm. Experiments were repeated in two independently differentiated batches. All data are expressed as mean ± SEM. Student’s t tests were performed to determine statistical significance. *p<0.05, **p<0.01, *** p<0.001, ****p<0.0001.

**Supplementary Table 1.**
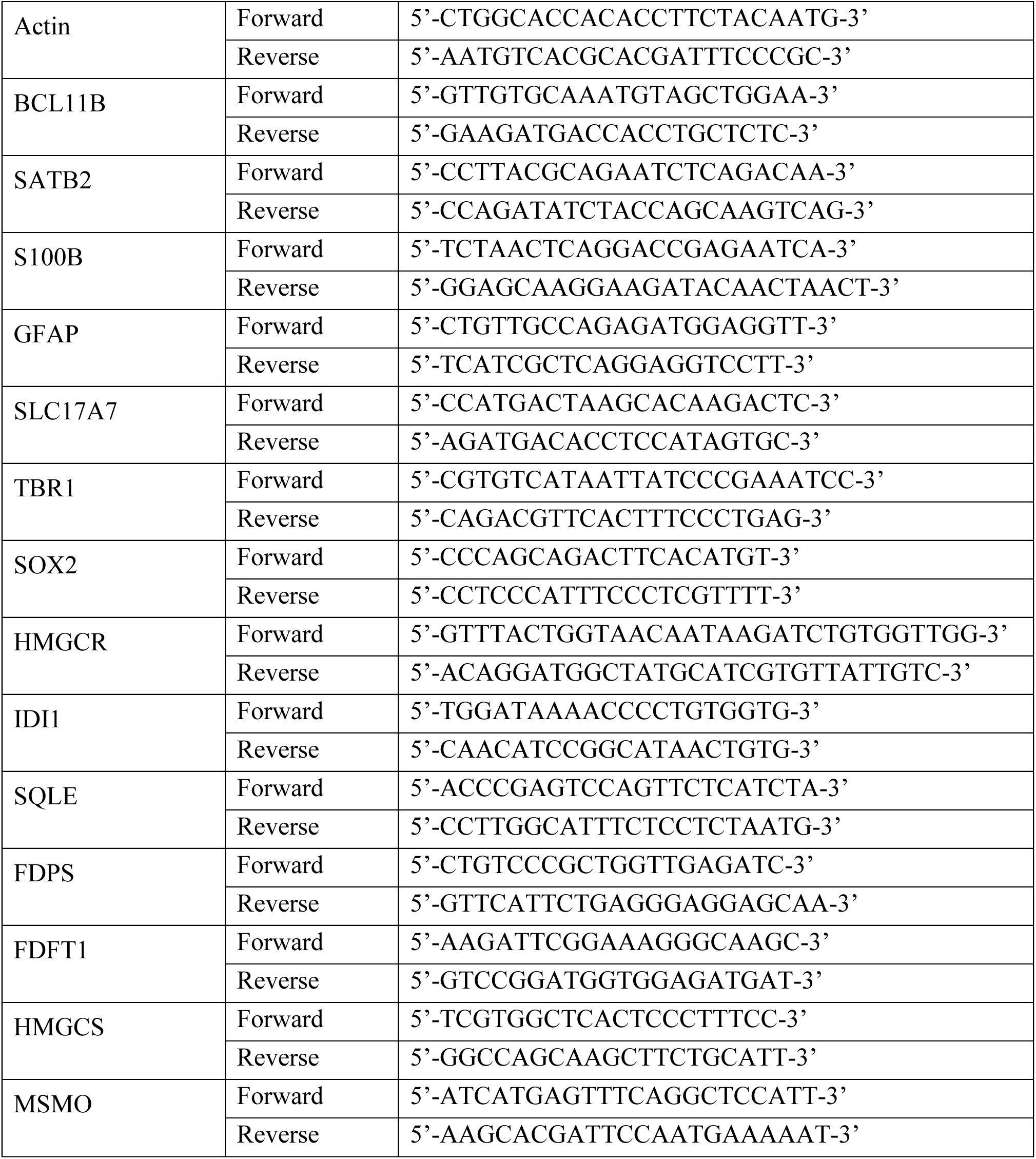
Primers information for RT-qPCR.

## Notes

### Competing Interest Statement

The authors have declared no competing interest.

